# Systematic Quantification of Sequence and Structural Determinants Controlling mRNA stability in Bacterial Operons

**DOI:** 10.1101/2020.07.22.216051

**Authors:** Daniel P. Cetnar, Howard M. Salis

**Affiliations:** Department of Chemical Engineering, The Pennsylvania State University, University Park, PA, 16802; Department of Biological Engineering, The Pennsylvania State University, University Park, PA, 16802; Department of Biomedical Engineering. The Pennsylvania State University, University Park, PA, 16802

## Abstract

mRNA degradation is a central process that affects all gene expression levels, and yet the determinants that control mRNA decay rates remain poorly characterized. Here, we applied a synthetic biology, learn-by-design approach to elucidate the sequence and structural determinants that control mRNA stability in bacterial operons. We designed, constructed, and characterized 82 operons, systematically varying RNAse binding site characteristics, translation initiation rates, and transcriptional terminator efficiencies in the 5’ UTR, intergenic, and 3’ UTR regions, and measuring their mRNA levels using RT-qPCR assays. We show that introducing long single-stranded RNA into 5’ UTRs reduced mRNA levels by up to 9.4-fold and that lowering translation rates reduced mRNA levels by up to 11.8-fold. We also found that RNAse binding sites in intergenic regions had much lower effects on mRNA levels. Surprisingly, changing transcriptional termination efficiency or introducing long single-stranded RNA into 3’ UTRs had no effect on upstream mRNA levels. From these measurements, we developed and validated biophysical models of ribosome protection and RNAse activity with excellent quantitative correspondence. We also formulated design rules to rationally control a mRNA’s stability, facilitating the automated design of engineered genetic systems with desired functionalities.

## INTRODUCTION

Engineering sophisticated genetic systems requires the development of more comprehensive biophysical models that can predict how changes to sequence affect gene expression levels, taking into account transcription, translation, and mRNA degradation (1–4). mRNA decay rates are an important contributor to genetic circuit function, altering the circuit’s dynamic and steadystate gene expression levels as well as controlling “turn off” times in response to changing transcriptional programs (5,6). While significant focus has been given to measuring and modeling transcriptional initiation rates (7–11), transcriptional control (12,13), and translation initiation rates (14–16), previous work on mRNA stability has primarily focused on discovering and characterizing the proteins and pathways responsible for mRNA decay (17–21).

A key challenge to modeling and predicting mRNA decay rates arises from the many participating enzymes and the high degree of coupling between transcription, translation, and RNAse activity (22,23). Predicting mRNA decay rates in multi-cistronic operons is further complicated by the potential for differential decay rates across coding sequences, called polarity (24,25). Here, we apply a learn-by-design approach to quantitatively measure how a bacterial mRNA’s sequence and structural properties determine its stability through the several mechanisms of mRNA decay. From these results, we formulate quantitative design rules for controlling a mRNA’s stability.

Several enzymes work together in a multi-pathway process to determine how quickly bacterial mRNAs are degraded (20,21), including endonucleases (*E. coli* RNAses E, G, III), exonucleases (*E. coli* RNAse II, PNPase, RNAse R), and the helper enzymes that modify the ends of mRNAs (*E. coli* RppH, DapF, PAPI) (**Figure 1**). These pathways are intimately coupled to both transcription and translation. Beginning with transcription initiation, the 5’ ends of all bacterial mRNAs contain a triphosphate group that inhibits RNAse activity. However, over time, RppH hydrolyzes these 5’ ends, yielding a monophosphate group that has higher affinity to RNAse E (26,27). RppH activity is promoted by the protein DapF and depends on the sequence and structure of at least the first three nucleotides of the transcript (**Figure 1A**) (28–30). As transcription elongation proceeds, co-transcriptional folding determines the types of mRNA structures that form and the portions that remain unstructured and accessible, which is also affected by RNA polymerases’ transcriptional elongation rate (31,32). During this time, as start codons are transcribed, ribosomes will bind and begin to translate the mRNA, following RNA polymerase and covering the mRNA with protective ribosomes. Once the mRNA is fully transcribed, the translation initiation and elongation rates of its coding sequences will determine the number of protective ribosomes bound to the transcript (**Figure 1B**).

**Figure 1:**
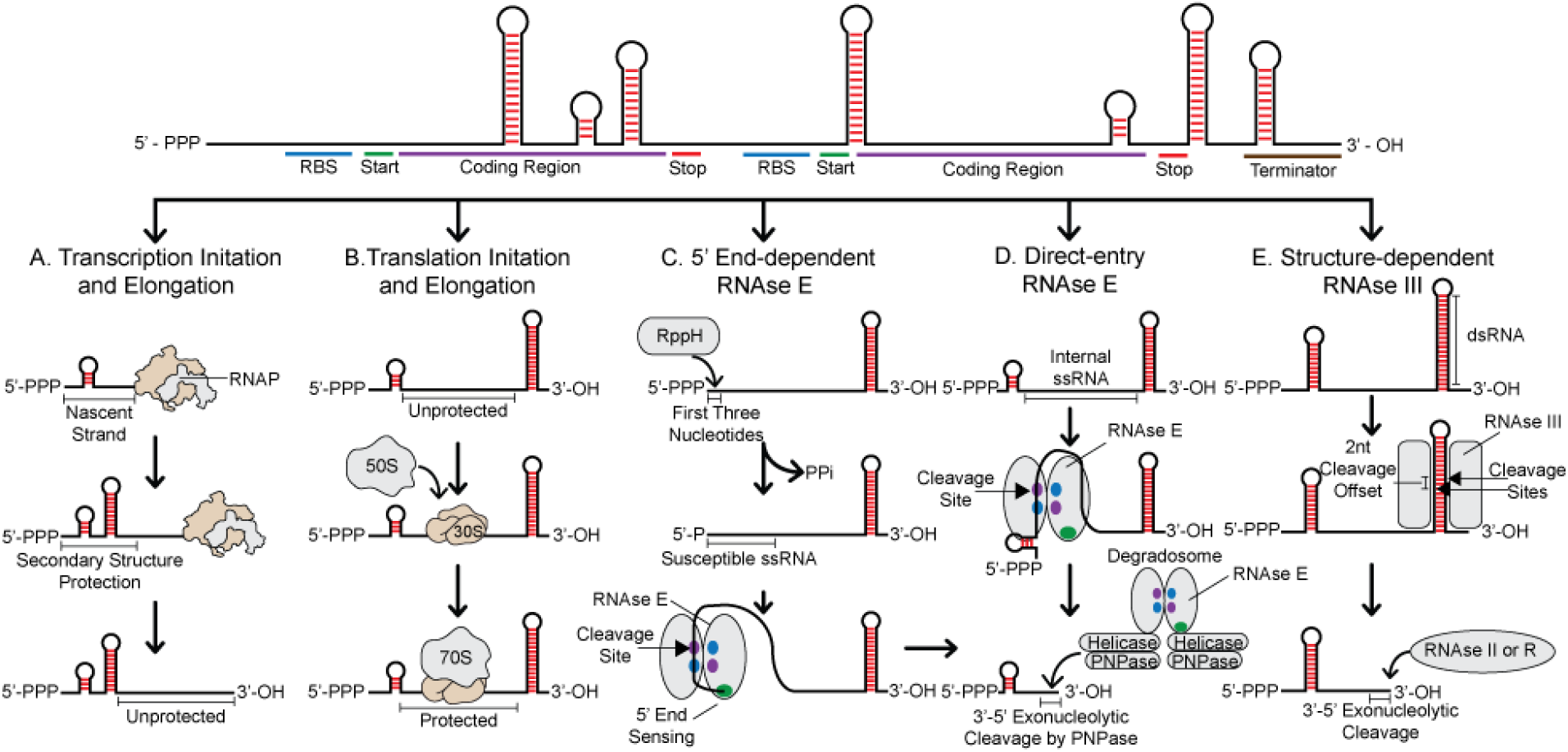
mRNA sequence determinants control mRNA Stability through coupling of transcription, translation, and RNAse activity. (A) Transcription elongation rates control mRNA structure and accessibility at 5’ ends. (B) Translation initiation and elongation rates control the number of protective ribosomes bound to mRNA. (C) Both mRNA structure and the first three nucleotides at mRNA 5’ ends controls RppH and RNAse E activity via the end-dependent decay pathway. (D) RNAse E binds and cleaves mRNA at single-stranded internal sites unprotected by structure. After initial cleavage, the RNA degradosome processively degrades mRNA. (E) RNAse III targets specific mRNA structures for cleavage. PNPase, RNAse II, and RNAse R degrade mRNA at 3’ ends using 3’ to 5’ exonuclease activity.

As transcription and translation proceeds, RNAses will interact with all accessible mRNA regions, binding and cleaving where possible. RNAse E and RNAse G cleave mRNAs at either their 5’ ends via end-dependent entry (**Figure 1C**) or at internal sites that remain accessible and unstructured (direct entry) (33–35) (**Figure 1D**). Other RNAses (e.g. RNAse III) bind to specific types of RNA structures, initiating their cleavage activity (36,37). Notably, an RNA endonuclease’s first cleavage event will create two new ends in the transcript, a monophosphate 5’ end and a 3’ end, that can then acted upon by the RNA degradosome (38), a loosely associated multi-protein complex, that combines both exonucleases and endonucleases (e.g. oligo-RNase, RNAse R, PNPase) to rapidly reduce the transcript to short oligonucleotides and mononucleotides (39,40). These activities include endonucleases that bind nearby monophosphate 5’ ends and cleave internally, releasing oligonucleotides (33) as well as exonucleases that bind to structurally accessible 3’ ends and chew them back in a 3’ to 5’ direction, releasing mononucleotides (41). Helper enzymes (e.g. PAPI) improve the accessibility of 3’ ends by adding unstructured polyA tails (39), accelerating exonuclease activity. Altogether, after the first cleavage event takes place, large regions of the mRNA transcript can be rapidly processed to mononucleotides. Overall, the rate of the first RNAse cleavage event is the slowest, making it a rate-limiting step in determining a mRNA’s overall stability (42). In particular, a first cleavage event inside the 5’ UTR or intergenic regions is often rapidly followed by destruction of start codons in nearby coding regions, eliminating protein expression.

Previous studies have qualitatively observed several factors that alter a mRNA’s decay rate, including (i) mRNA sequence motifs that may specifically bind RNAses (43,44); (ii) specific sequences at a mRNA’s 5’ end that alter RppH binding (29,45); (iii) the co-transcriptional, temperature-sensitive formation of mRNA structures that block RNAse binding and increase mRNA stability (46); (iv) the binding of ribosomes in 5’ UTRs and coding regions that block RNAse binding, called protective ribosomes (24,47,48); (v) RNA structures at transcript 3’ ends that block exonuclease activities (49); and (vi) changes in environmental growth conditions that differentially alter RNAse and helper enzyme levels (e.g. stress responses) (50). RNAse III is also known to cleave long RNA duplexes (51,52), while specific motifs, such as RAUUW (44), RNWUU (53), and RNAU (33,54), were suggested to specifically bind RNAse E. Overall, *E. coli* natural mRNAs have half-lives of about 1 to 10 minutes with the potential to vary protein expression levels by at least 10-fold (31,55).

However, the precise sequence determinants that alter mRNA stability have not been quantitatively elucidated. Prior studies suggesting specific binding motifs have relied on a small number of characterized mRNAs, while the relationship between RppH activity, translation rates, transcriptional termination efficiency, and transcript stability remains poorly characterized. Because natural mRNAs are often attacked by RNAses at multiple entry sites, it remains challenging to interpret natural sequence-stability assays and identify quantitative cause-effect relationships. Instead, we use a synthetic biology, learn-by-design approach to systematically alter a mRNA transcript’s properties and measure its mRNA levels during sustained exponential growth conditions. Through this strategy, we designed, constructed, and characterized the mRNA levels from 82 mono- and bi-cistronic operons, systematically varying the mRNA’s sequence and structural properties to modulate RppH, RNAse E/G, and RNAse III activities. We also systematically varied the mRNA’s translation rates and transcriptional termination efficiencies to quantitatively determine how they protect mRNA transcripts from endonuclease or exonuclease activities. Altogether, our dataset provides a quantitative and comprehensive understanding of how mRNA sequence controls mRNA stability through several interactions.

## MATERIAL AND METHODS

### Plasmid Design and Cloning

A series of libraries of pFTV1-derived plasmids with a ColE1 origin of replication and Chloramphenicol resistance (Cm^R^) were designed and constructed to express fluorescent protein reporters with the objective of testing specific design motifs’ effect on mRNA stability. In **Supplementary Data 1**, the dataset name and sequence information are listed for each construct. For the datasets mRFP1 Expression Library and Terminator Efficiency, a base monocistronic mRFP1 pFTV1-derived plasmid was used that contained the σ^70^ constitutive promoter J23100 and a double terminator. To create the mRFP1 Expression Library, the 5’ UTR was engineered using RBS Library Calculator to design a set of ribosome binding sites that varied the expression of mRFP1 1000-fold. The degenerate oligonucleotide (IDT) designed by RBS Calculator containing these designs was flanked by a 5’ XbaI and 3’ NdeI restriction site and primer binding sites, which allowed for PCR amplification. The plasmid was then assembled using cut and paste cloning. Similarly, the Terminator Efficiency set used construct 6 from the mRFP1 Expression Library as the base plasmid. The terminators tested were assembled by annealing two complementary oligonucleotides (IDT) with 5’ FseI and 3’ SphI restriction overhangs. The annealed oligonucleotides were ligated into the base plasmid replacing the existing terminators. Likewise, for the datasets sfGFP Expression Library and sfGFP Coupled Expression, the base plasmid contained the σ^70^ constitutive promoter J23100 and a double terminator. RBS Libraries were designed and cloned in for both using a similar method as for the mRFP1 Expression Library, except that the restriction sites employed were 5’ XbaI and 3’ AgeI. For the datasets, 5’ Variable Length Poly A, Intergenic Variable Length Poly A, RNAse Polarity, and RppH Set, a base bi-cistronic codon optimized mRFP1 and GFPmut3B pFTV1-derived plasmid was used that contained the σ^70^ constitutive promoter J23100 and a double terminator. At the 5’ UTR, RBS Calculator v2.1 was used to design a ribosome binding site sequence with an XbaI restriction site on the 5’ end and with a translation initiation rate of 9395 au on RBS Calculator’s proportional scale. The newly designed 5’ UTR and J23100 promoter was assembled by annealing two complementary oligonucleotides (IDT) with 5’ BamHI and 3’ SacI restriction overhangs and ligating them into the base plasmid. Likewise, RBS Calculator v2.1 was used to design a ribosome binding site sequence with an AatII restriction site on the 5’ end with a translation initiation rate of 7083 au on RBS Calculator’s proportional scale. The newly designed intergenic region was assembled by annealing two complementary oligonucleotides (IDT) with 5’ EcoRI and 3’ XhoI restriction overhangs and ligating them into the base plasmid. Additions to the base 5’ UTR were made using a 5’ BamHI site upstream of the promoter and the newly added 3’ XbaI site. Sequence additions to the base intergenic UTR was made using an upstream 5’ EcoRI site and the newly added 3’ AatII site. For additions to the 3’ UTR, a 5’ PacI site and 3’ SpeI site were used. All additions were made by annealing complementary oligonucleotides (IDT) with the correct restriction overhangs for the 5’ UTR, intergenic region, and 3’ UTR. All plasmids were transformed into *Escherichia coli* DH10B cells, followed by sequence verification of isolated clones.

### Strains, Growth and Characterization

All measurements were conducted using *Escherichia coli* DH10B cells containing plasmids, cultured using M9 minimal media, and maintained in the exponential growth phase for at least 20 hours. For each construct, isogenic colonies were used to inoculate overnight cultures in 500 ul LB media supplemented with 50 ug/mL Chloramphenicol (Cm) in a 96-well deep-well plate. Overnight cultures were diluted 100-fold by diluting 2 uL of culture into 198 ul of M9 minimal media with Cm using a 96-well microtiter plate, and incubated at 37 °C with high orbital shaking inside a Spark spectrophotometer (TECAN). OD_600_ absorbance was taken every 10 minutes until the OD_600_ reached 0.15-.20, indicating the cells’ entry into the mid-exponential phase of growth. At this time, a subsequent 96-well microtiter plate was prepared by serial dilution of the culture from the first plate into M9 minimal media with Cm maintaining the cells in the exponential phase of growth. For each culture, single-cell red florescent protein (RFP) or green florescent protein (GFP) fluorescence measurements were performed by collecting 10 ul from the end of the second dilution, transferring to a microtiter plate with 200 ul PBS solution with 2 mg/ml kanamycin, and recording 100,000 single-cell fluorescence levels with a Fortessa flow cytometer (BD Biosciences). All single-cell fluorescence distributions were unimodal. The mean of the distributions is calculated, and the background autofluorescence of *Escherichia coli* DH10B cells subtracted. All reported fluorescence levels are the average of at least two biological replicates, which are listed in **Supplementary Data 1**.

mRNA level measurements were performed on selected strains by inoculating a 5 mL culture of Cm supplemented LB media and incubated at 37 °C with 300 RPM shaking. Once cells reached an OD600 of 1.0, measured using a cuvette-based spectrophotometer (NanoDrop 2000C), they were diluted to an OD_600_ of 0.05 in a 5 mL culture of Cm supplemented M9 minimal media and incubated at 37 °C with 300 RPM shaking. Once cells reached an OD_600_ of between 0.4-0.6, they were diluted to OD 0.001 in a 5 mL culture of Cm supplemented M9 minimal media and incubated at 37 °C with 300 RPM shaking. Cells were harvested once they reached an OD_600_ of between 0.15-0.25 and their total RNA extracted using the Total RNA Purification kit (Norgen Biotek), followed by non-specific degradation of contaminant DNA using the Turbo DNAse kit (Ambion). Following extraction, cDNA was prepared using the High Capacity cDNA Reverse Transcription kit (Applied Biosystems). Taqman based qPCR was performed using an ABI Step One Plus real-time thermocycler (Applied Biosystems), utilizing a Taqman probe targeting the constructs of interest mRFP1, codon optimized mRFP1, sfGFP, codon optimized GFPmut3B. Additionally, a custom 16S rRNA TaqMan probe was used as an endogenous control and was used to calculate relative mRNA levels from ΔC_t_ numbers. TaqMan probes sequence and validation are listed in Supplementary Data 2. Likewise, SYBR Green based qPCR was performed in a similar manner for the determination of the relative mRNA level.

## RESULTS

### Unstructured RNA in the 5’ Untranslated Region Controls mRNA Stability

We first interrogated the sequence and structural determinants in the 5’ untranslated region that affect a mRNA’s stability in *E. coli*, combining systematic design, DNA assembly, and characterization using RT-qPCR and fluorescence measurements. To do this, we designed and constructed plasmid-encoded, bi-cistronic operons that utilize a constitutive sigma70 promoter (J23100), a rationally designed synthetic ribosome binding site, and two codon-optimized fluorescent protein reporters (mRFP1 and GFPmut3) (**Methods**, **Figure 2A**). The ribosome binding sites were designed by the RBS Calculator v2.1 to have moderate translation initiation rates (7000 to 10000 au). Using a sequence constraint, the RBS designs included a fast-folding, insulating mRNA hairpin (ΔG_folding_ = −12.61 kcal/mol) that prevents upstream sequence changes from changing the mRNA’s translation initiation rate. Rationally designed upstream sequences were then inserted to replace the 5’ untranslated region, beginning with the transcriptional start site and ending past the coding region’s start codon. We then carried out long-time cultures of transformed strains, maintaining them in the exponential growth phase with periodic serial dilutions, followed by mRNA level measurements using RT-qPCR assays with TaqMan probes specific to the internal regions of reporter coding sequences (**Methods**). All sequences and measurements are located in **Supplementary Data 1**. All TaqMan probe sequences and measured efficiencies are located in **Supplementary Data 2**. The MIQE information for these experiments is located in **Supplementary Data 3**. By keeping the promoter unchanged and maintaining cells in the exponential growth phase for a long time period, changes in mRNA level can be primarily attributed to changes in mRNA stability, although we discuss potentially confounding factors.

**Figure 2:**
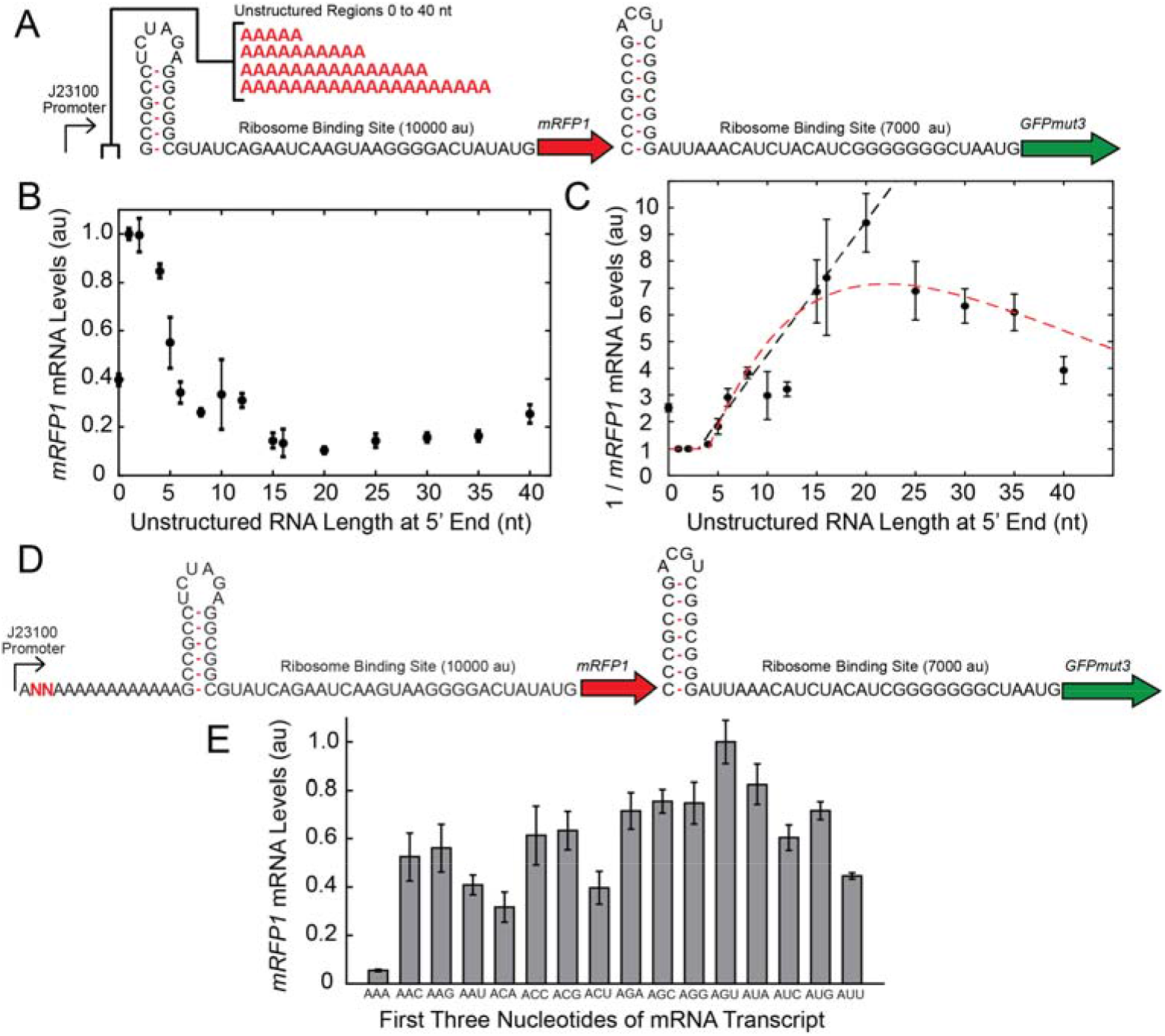
Sequence determinants of mRNA stability at 5’ ends. (A) Schematic showing a mRFP1-sfGFP operon with systematically varied 5’ UTR sequences with between 0 to 40 nucleotides of single-stranded RNA, inserted upstream of an insulating hairpin. (B) RT-qPCR measurements of *mRFP1* mRNA levels show a quantitative relationship between single-stranded RNA length and mRNA stability. Dots and error bars are the mean and standard deviation of 2 biological and 6 technical replicates. (C) RT-qPCR measurements are replotted showing the relationship between the inverse mRNA level (proportional to decay rate) and single-stranded RNA length. Dashed black and red lines show predictions from two biophysical models. (D) Schematic showing a mRFP1-sfGFP operon with systematically varied dinucleotides at the 5’ end. (E) RT-qPCR measurements of *mRFP1* RNA levels show how varied dinucleotides affected mRNA stability. Bar heights and error bars are the mean and standard deviation of 2 biological and 6 technical replicates.

In the first series of 16 operons, we designed and inserted 5’ UTR sequences that contained between 0 to 40 polyA nucleotides, creating systematically longer single-stranded RNA (ssRNA) regions upstream of the ribosome binding site’s insulating hairpin (**Figure 2A**). Using Vienna RNAfold (56), we verified that these polymeric A sequences do not fold into mRNA structures that could potentially block RNAse endonuclease activity. After characterizing these operons, we found a clear, quantitative relationship between the length of the ssRNA region and the resulting *mRFP1* mRNA level (**Figure 2B**). The highest mRNA levels occurred when the 5’ UTR contained very short polyA sequences (2 or 3 As), followed by a precipitous decrease as the ssRNA region was lengthened. mRNA levels were reduced by about 2-fold when a ssRNA region was 5 nucleotides long. At its lowest, mRNA levels were reduced by 9.4-fold when 20 ssRNA nucleotides were added to the 5’ UTR. However, further lengthening of the ssRNA region reversed the trend; we found that mRNA levels consistently increased when the ssRNA region was 20 to 40 nucleotides long. Surprisingly, without a polymeric A region (0 As), the presence of an insulating hairpin at the transcriptional start site actually reduced mRNA levels by about 2.5-fold.

With further analysis, we then developed biophysical models to explain how lengthening the ssRNA region could alter mRNA levels in such a non-linear fashion. When cultures are maintained in the exponential phase of growth, mRNA levels will be inversely proportional to their decay or degradation rates. We therefore replotted our data to show how the inverse of mRNA levels are related to the lengths of the ssRNA region (**Figure 2C**). The trends in this relationship immediately suggested two types of biophysical models. In both models, RNAses E/G bind and cleave mRNA anywhere where is a minimally sized landing pad containing an unstructured ssRNA region. If the unstructured region is too short, RNAses E/G cannot bind and cleave mRNA. As the unstructured region is lengthened, the number of potential binding sites also increases, resulting in a proportional increase in RNAse activity.

In the first model, we calculate the change in a mRNA’s decay rate Δδ_mRNA_ by only considering the length of the smallest possible binding site for RNAses E/G (N), the length of a ssRNA region (L), and an activity coefficient (C), according to the equation:

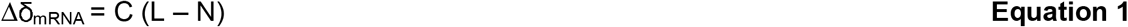

With a minimum binding site of 2 nucleotides (N = 2 nt) and an activity coefficient of about one half (C = 0.528 1 / nt), Equation 1 can explain how mRNA decay rates depend on ssRNA regions from 1 to 20 nucleotides long (Pearson R^2^ = 0.92, mean absolute error = 0.473) (**Figure 2C**, black dashed line). However, it’s clear that this linear relationship is no longer true after the ssRNA region is more than 20 nucleotides long.

In the second model, we additionally take into account that untethered ssRNA molecules can dynamically form transient loop structures that limit their accessibility, and therefore may prevent RNAses from binding to them. From the wormlike chain model of polymer theory, the likelihood that ssRNA with length L will *not* form an intramolecular loop is proportional to exp(−L / P), where P is the persistence length of ssRNA, which is a measure of its bending stiffness. As the length of a ssRNA region exceeds the persistence length, it becomes highly likely that the ssRNA will fold into a disordered structure. We therefore incorporated this effect to calculate the change in a mRNA’s decay rate according to the equation:

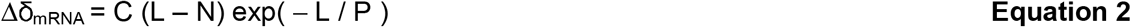

The persistence length of a ssRNA region depends on several properties, including RNA sequence, temperature, and the solution’s salt composition, with previous measurements ranging from 1 to 7 nm or roughly 3 to 21 nucleotides (57–60). Here, with a minimum binding site of 2 nucleotides (N = 2 nt), an activity coefficient of about one (C = 0.975 1/nt), and a persistence length of 21 (P = 21 nt), we found that Equation 2 can explain how the mRNA’s decay rate depended on ssRNA region’s length from 1 to 40 nucleotides (Pearson R^2^ = 0.874, mean absolute error = 0.767) (**Figure 2C**, red dashed line). Overall, the second model better predicts these mRNAs’ decay rates across the entire dataset. However, when there is no singlestranded region (L = 0 nt), we do observe a discontinuity in both model predictions that could be caused by a confounding effect by another interaction, for example, a reduction in transcription rate.

### First Three Nucleotides in the 5’ Untranslated Region Control a mRNA’s Stability

Previous *in vitro* measurements have shown that the first three nucleotides of a mRNA transcript have an effect on the rate of 5’ end dephosphorylation by RppH or RppH-DapF complex (26), which will affect RNAse E/G’s ability to initiate mRNA decay via the end-dependent pathway. Here, we carried out *in vivo* measurements to quantify how the first three nucleotides of a mRNA transcript altered its mRNA level. We designed and constructed a series of 16 bi-cistronic operons that express mRFP1 and GFPmut3 where the 5’ UTR contains a ‘ANN’ 5’ end, followed by a polymeric 15A region, an insulating RNA hairpin, and a designed ribosome binding site with a translation initiation rate of about 10000 au on the RBS Calculator v2.1 scale (**Figure 2D**). The polyA region prevents the formation of mRNA structures regardless of the first three nucleotides in the transcript, which was verified by using Vienna RNAfold (56). After maintaining cell cultures in the exponential growth phase, we carried out RNA extractions and RT-qPCR to measure *mRFP1* mRNA levels. We found that *mRFP1* mRNA levels varied by up to 18.2-fold when varying just the first three nucleotides of the transcript with the 5’-AAA-3’ and 5’-AGU-3’ trinucleotides have the most and least effect, respectively (**Figure 2E**). Even when excluding the most potent variant 5’-AAA-3’, mRNA levels still varied by 3.2-fold, which is a potent effect for such a small sequence region. These results provide further support that the 5’ end sequence affects how RppH and DapF interact with mRNA to accelerate end-dependent mRNA decay.

### mRNA Translation Initiation Rate Controls mRNA Stability via Ribosome Protection

Previous studies have suggested that ribosomes can protect mRNA transcripts from RNAse activity, possibly by covering RNAse binding sites and limiting their accessibility (61,62) (**Figure 3A**). However, the relationship between a mRNA’s translation rate and its decay rate has never been quantitatively studied. To determine such a quantitative relationship, we designed and constructed 18 mono-cistronic operons expressing either mRFP1 or sfGFP reporter, systematically varying their translation initiation rates by up to 1300-fold (**Figure 3B**). To do this, we inserted optimized RBS libraries into their 5’ UTRs, each designed by the RBS Library Calculator to have an insulating hairpin, followed by a small number of pinpoint mutations that would greatly change the reporters’ translation initiation rates (63). We then characterized the operons’ expression levels, measuring both their mRNA and protein levels using RT-qPCR and flow cytometry, respectively (**Methods**).

**Figure 3:**
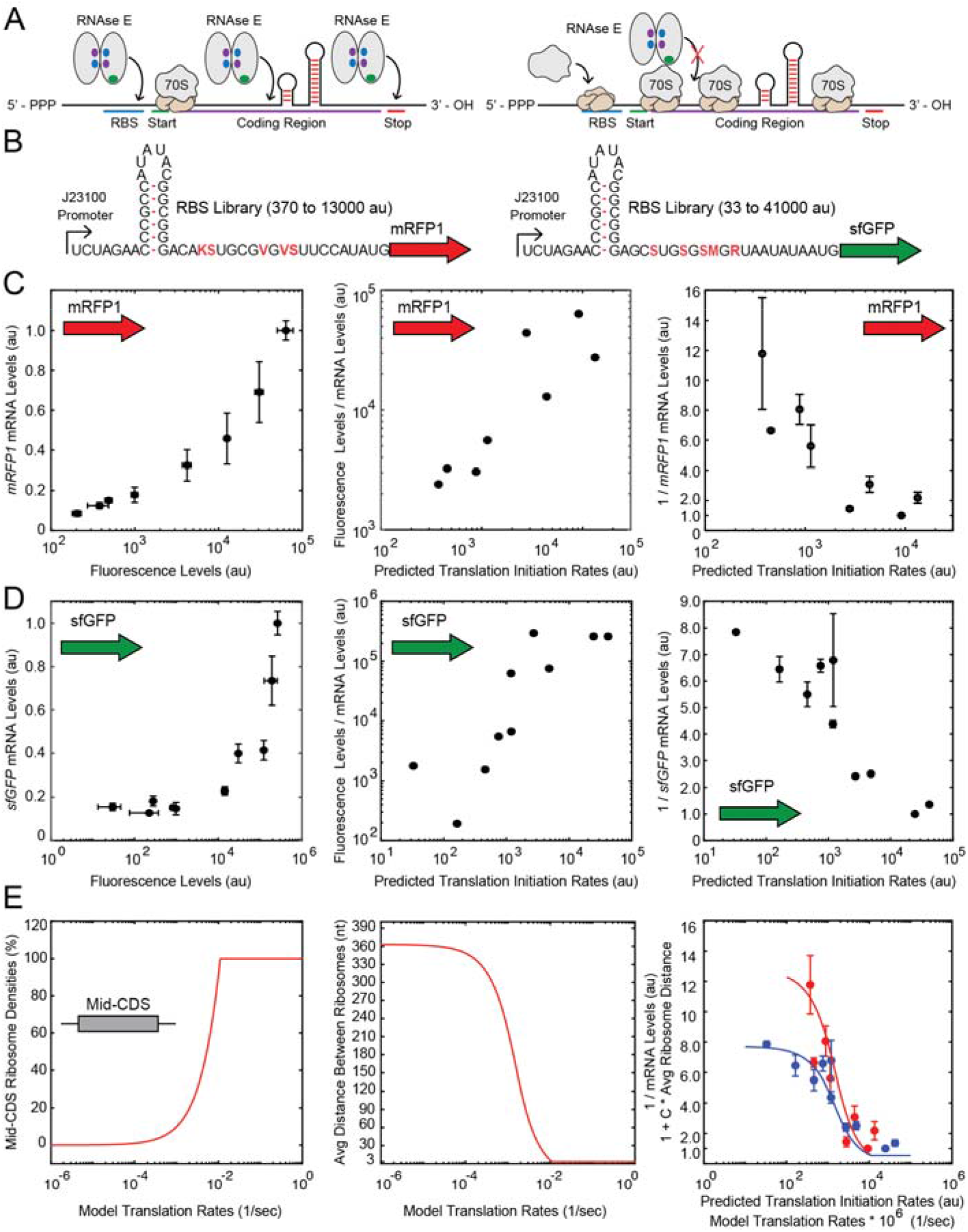
Translation rate and mRNA stability are coupled via ribosome protection. (A) Schematics illustrating how a mRNA’s translation rate controls the accessibility of RNAse binding sites in an operon’s coding regions. (B) Schematics showing the RBS library designs inside operons expressing either *mRFP1* or *sfGFP* reporter proteins. Highlighted nucleotides indicate the pinpoint mutations in RBS libraries. (C) Characterization of the *mRFP1* operons, showing (left) *mRFP1* mRNA levels compared to mRFP1 fluorescence levels, (middle) the apparent mRFP1 translaton rates (the ratio of mRFP1 fluorescence over *mRFP1* mRNA levels) compared to the *mRFP1* predicted translation initiation rates, and (right) the inverse *mRFP1* mRNA levels compared to the *mRFP1* predicted translation initiation rates. (D) Characterization of the *sfGFP* operons, showing (left) *sfGFP* mRNA levels compared to sfGFP fluorescence levels, (middle) the apparent *sfGFP* translation rates (the ratio of sfGFP fluorescence over *sfGFP* mRNA levels) compared to the *sfGFP* predicted translation initiation rates, and (right) the inverse *sfGFP* mRNA levels compared to the *sfGFP* predicted translation initiation rates. (E) Calculations from a biophysical model of ribosome protection show (left) how translation rate controls ribosome density, (middle) how translation rate controls the average distance between adjacent ribosomes, and (right) a comparison between the average distance between ribosomes and the characterized mRNA decay rates (inverse mRNA levels) for (red) *mRFP1* and (blue) *sfGFP*. Dots and error bars showing mRNA level measurements are the mean and standard deviation from 2 biological and 6 technical replicates. Dots and error bars showing fluorescence level measurements are the mean and standard deviation from 2 to 4 biological replicates. Predicted translation initiation rates use the RBS Calculator v2.1 proportional scale.

We found that systematically increasing the mRNAs’ translation initiation rates resulted in corresponding increases in both their mRNA and protein levels. For example, increasing the *mRFP1* translation initiation rate from 370 to 13000 au on the RBS Calculator v2.1 scale resulted in a 11.8-fold increase in *mRFP1* mRNA level and a 313-fold increase in mRFP1 fluorescence level (**Figure 3C**, left). Likewise, increasing the sfGFP translation initiation rate from 33 to 41000 au resulted in a 7.85-fold increase in sfGFP mRNA level and a 8760-fold increase in sfGFP fluorescence level (**Figure 3D**, left).

Here, the higher protein levels originated from both higher translation initiation rates and higher mRNA levels. To distinguish these sources, we calculated the apparent translation rates by taking the ratio of fluorescence level over mRNA level, and compared them to the RBS Calculator’s predicted translation initiation rates (**Figure 3CD**, middle). We confirmed excellent quantitative agreement between the predicted and measured translation rates for both *mRFP1* and *sfGFP* with strong statistical significance (*mRFP1*: Pearson R^2^ = 0.782, *p* = 0.0035; *sfGFP*: Pearson R^2^ = 0.706, *p* = 0.0023). We then replotted the same predictions and measurements to determine how the mRNAs’ decay rates – the inverse of our mRNA level measurements – were affected by the mRNA’s translation initiation rates (**Figure 3CD**, right). As expected, the lowest translation initiation rates yielded the highest mRNA decay rates, qualitatively consistent with a ribosome protection mechanism. However, quantitatively, our measurements could equally support a log-linear or sigmoidal relationship between translation rate and mRNA decay. To investigate further, we developed a biophysical model of ribosome protection that provides support for a sigmoidal relationship.

### A Biophysical Model of Ribosome Protection Explains mRNA Stability

Consider a mRNA transcript that contains a protein coding sequence (CDS) with *L_mRNA_* amino acids. The key question is to calculate how many of these nucleotides remain unprotected by elongating ribosomes. Here, we designate α as the ratio of the ribosome’s translation initiation rate over its elongation rate. We also designate *F* as the physical footprint of each ribosome in units of trinucleotides (amino acids). Based on prior measurements, we specify that the ribosome’s footprint *F* is about 10 trinucleotides (30 nt). According to a TASEP (totally asymmetric exclusion process) model of ribosome dynamics that includes the ribosome’s footprint on the mRNA (64), designated as an “extended object”, a single equation can be used to approximately calculate how the translation initiation rate controls the ribosome’s average density. When translation initiation is the rate-limiting step (α is less than one), the steady-state ribosome density *ρ_r_* is:

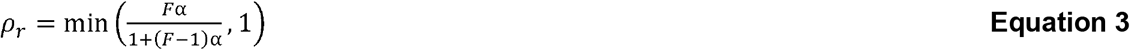

when averaged over the middle portion of the CDS, away from the start and stop codons. The density of protected mRNA is *Fρ_r_*, which includes the ribosome’s footprint. By definition, the maximum number of bound ribosomes is *L_mRNA_/F* with a density of one. The relationship defined by **Equation 3** illustrates how increasing a mRNA’s translation initiation rate results in higher ribosome densities, up until the maximum possible value (**Figure 3E**, left).

We next consider the distances *D* between adjacent ribosomes, called the “headway distance” from studies of vehicular traffic. Here, the absence of a bound ribosome at a position along the mRNA is called a “hole”. From the TASEP model with extended objects (65), the probability that a bound ribosome has *m* free trinucleotides in front of it is related to both the ribosome density *ρ_r_* and hole density *ρ_h_* according to:

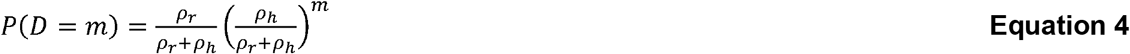

where the hole density is determined by *ρ_h_* = 1 – *Fρ_r_*. The first part of **Equation 4** calculates the probability that a mRNA position is bound by a ribosome whereas the second part calculates the probability that the next *m* adjacent positions on the mRNA all contain a hole. We then substitute **Equation 3** and the definition of the hole density into **Equation 4**, yielding the probability distribution for the ribosomes’ headway distances in terms of the ribosome density and footprint:

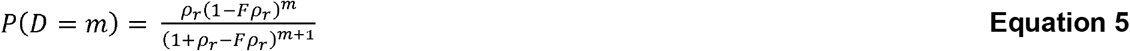

We then determine the ribosomes’ average headway distance by calculating the first moment of the probability distribution in **Equation 5**, using:

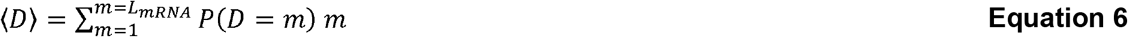

Altogether, for each selected translation initiation rate, we calculate the ribosome density using **Equation 3**, substitute it into **Equation 5** across a range of potential distances, and use this distribution and **Equation 6** to calculate the average distance between ribosomes (**Figure 3E**, middle). The overall relationship is a sigmoidal curve; at the highest translation initiation rates, the average distance plateaus to the smallest possible value (1 trinucleotide), whereas at the lowest translation initiation rates, the average distance plateaus at about *L_mRNA_*/2.

Finally, according to a ribosome protection mechanism, we anticipated that the average distance between ribosomes – the amount of unprotected mRNA – is proportional to the mRNAs’ decay rates. We therefore compared our inverse mRNA level measurements to the average distances between ribosomes across the 18 operons expressing *mRFP1* or *sfGFP* at varying translation initiation rates (**Figure 3E**, right) and found strong quantitative agreement with statistical significance (Pearson R^2^ = 0.823, *p* < 10^-6^), using proportionality constants C_mRFP1_ = 0.10 and C_sfGFP_ = 0.058. Here, the proportionality constant combines several factors, including the RNAse E/G activity per unprotected mRNA, the likelihood that unprotected mRNA forms protective structures, and the scale of the mRNA level measurements. The developed biophysical model of ribosome protection shows that the relationship between a mRNA’s translation initiation rate and its decay rate is expected to be a sigmoidal curve, which is well supported by our measurements.

### Location of RNAse Binding Sites Controls mRNA Stability in Bicistronic Operons

We next investigated how the locations of RNAse binding sites – inside either the 5’ UTR, intergenic region, or 3’ UTR regions – affected the mRNA levels of individual cistrons within a bicistronic operon. To do this, we designed and constructed 6 bicistronic operons expressing *mRFP1* and *GFPmut3*, inserting either a 20A ssRNA region (an RNAse E/G site) or a 25 bp RNA hairpin (a RNAse III site) into these three separate locations (**Figure 4A**). We applied Vienna RNA folding calculations to verify that the 20A region does not fold into a mRNA structure after being inserted into these locations. We performed similar calculations to verify that the 25 bp RNA hairpin does not fold into an alternative structure when inserted. We also constructed baseline operons that did not contain inserted RNAse binding sites, which we used as the reference for comparisons. Otherwise, all operons utilized the same promoters, ribosome binding sites, and coding regions. Based on prior studies of gene expression polarity (49,66–68), we anticipated the possibility that *mRFP1* and *GFPmut3* mRNA levels could be differentially affected by these inserted RNAse binding sites.

**Figure 4:**
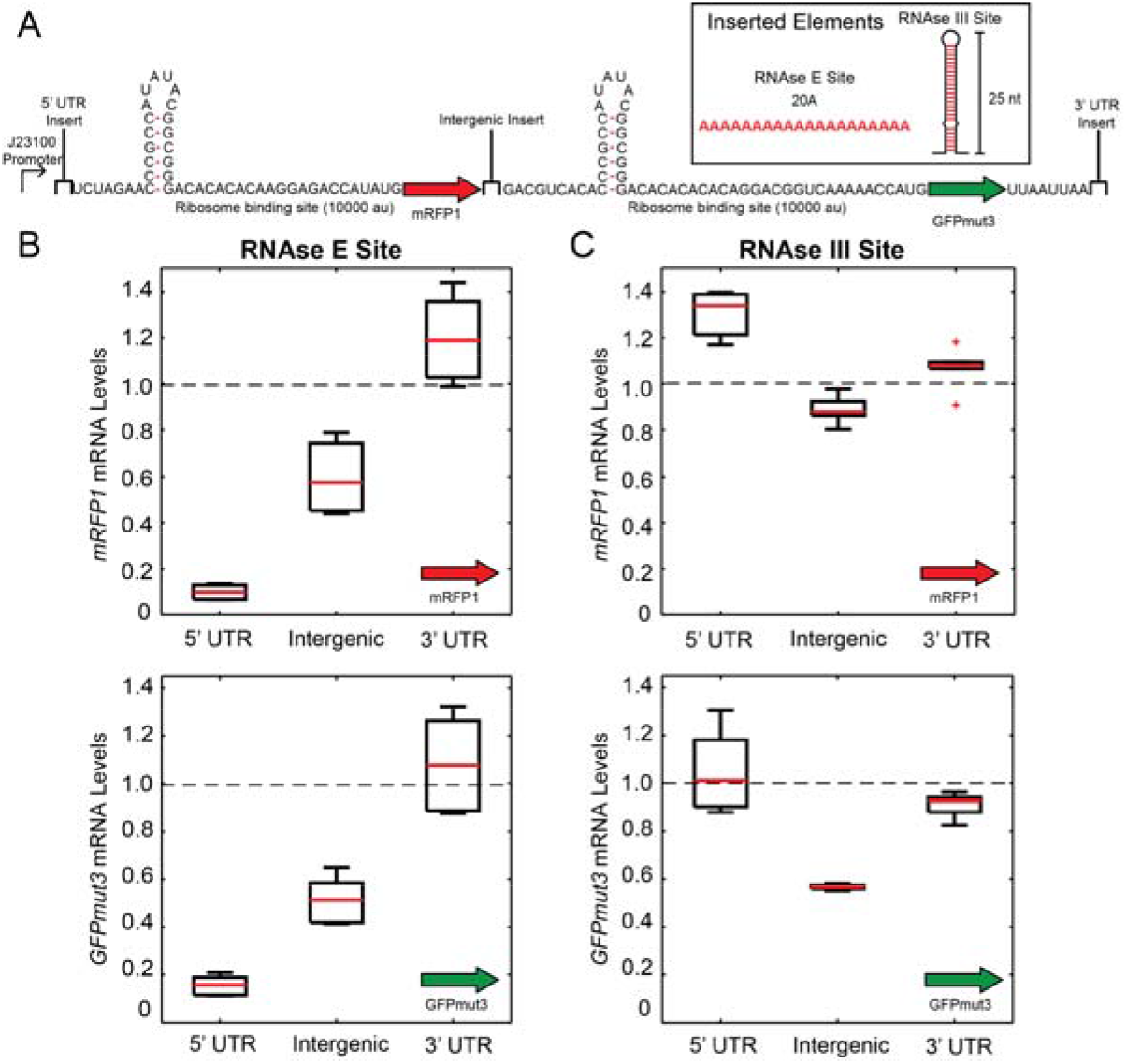
Positions of RNAse sites control mRNA stability. (A) Schematic illustrating the bistronic operons and the locations where RNAse E/G or RNAse III sites were inserted. (boxed) The two types of inserted RNAse binding sites are shown: either a 20A single-stranded RNA site or a 25 bp RNA hairpin structure. (B) Characterization of the operons, showing how the positions of inserted RNAse E sites affected the *mRFP1* and *sfGFP* mRNA levels. (C) Characterization of the operons, showing how the positions of inserted RNAse III sites affected the *mRFP1* and *sfGFP* mRNA levels. Dots and error bars are the mean and standard deviation from 2 biological and 6 technical replicates.

As before, we characterized these operons by extracting total RNA from cells maintained in the exponential growth phase and applying RT-qPCR assays to separately measure *mRFP1* and *GFPmut3* mRNA levels (**Methods**). Consistent with our prior results (**Figure 2B**), we found that inserting a 20A ssRNA region into the 5’ UTR greatly reduced the *mRFP1* mRNA levels (by 11fold) with a concomitant decrease in *GFPmut3* mRNA levels (by 7-fold) (**Figure 4B**). However, inserting the same 20A ssRNA region into the intergenic region had a much lower effect; *mRFP1* and *GFPmut3* mRNA levels were reduced by only 1.7-fold and 2-fold, respectively. Notably, inserting the 20A ssRNA region into the 3’ UTR region had no appreciable effect on the mRNA levels of either cistron. Overall, we found that RNAse E/G sites had a potent effect on a mRNA’s stability with a clear position-dependent trend; upstream sites accelerated mRNA decay more so than downstream sites. However, we did not observe any polarity effects; each RNAse E/G site similarly affected the mRNA levels of the operon’s *mRFP1* and *GFPmut* cistrons.

In contrast, only one of the inserted RNAse III sites had an appreciable effect on the operons’ mRNA levels, though it was accompanied by evidence of polarity (**Figure 4C**). Inserting a 25 bp RNA hairpin into the intergenic region did not appreciably affect the *mRFP1* mRNA level (1.13-fold change), but did decrease the *GFPmut3* mRNA level by 1.8-fold. Inserting the same 25 bp RNA hairpin into either the 5’ UTR or 3’ UTR had no appreciable effect on either mRNA level. Overall, these results suggest that the directionality and processivity of mRNA decay depend on the RNAse that carries out the first cleavage event.

### Unstructured RNA in the Intergenic Region Modestly Affects mRNA Stability

Following the previous results, we carried out a systematic investigation to quantitatively determine how single-stranded RNA (ssRNA) sites inside the intergenic region control mRNA stability. We designed and constructed 16 bicistronic operons expressing *mRFP1* and *GFPmut3*, inserting polyA ssRNA sites inside their untranslated intergenic regions, ranging from 0 to 40 nt long (**Figure 5A**). We then carried out RNA extractions on cells maintained in the exponential growth phase, and measured both *mRFP1* and *GFPmut3* mRNA levels using RT-qPCR assays. Consistent with our previous results, we found that unstructured RNA inside the intergenic region has an appreciable, but modest, effect on mRNA stability, affecting both upstream and downstream cistrons similarly (**Figure 5B**). When the ssRNA length is small (0 to 2 nt long), there is small rising trend in *mRFP1* and *GFPmut3* mRNA levels, reaching up to a 1.4-fold increase at their maximum. However, further increases in ssRNA length (3 to 40 nt) reversed the trend; both mRNA levels decreased by modest amounts, reaching a minimum of 1.66-fold lower *mRFP1* and 1.43-fold lower *GFPmut3* mRNA levels.

**Figure 5:**
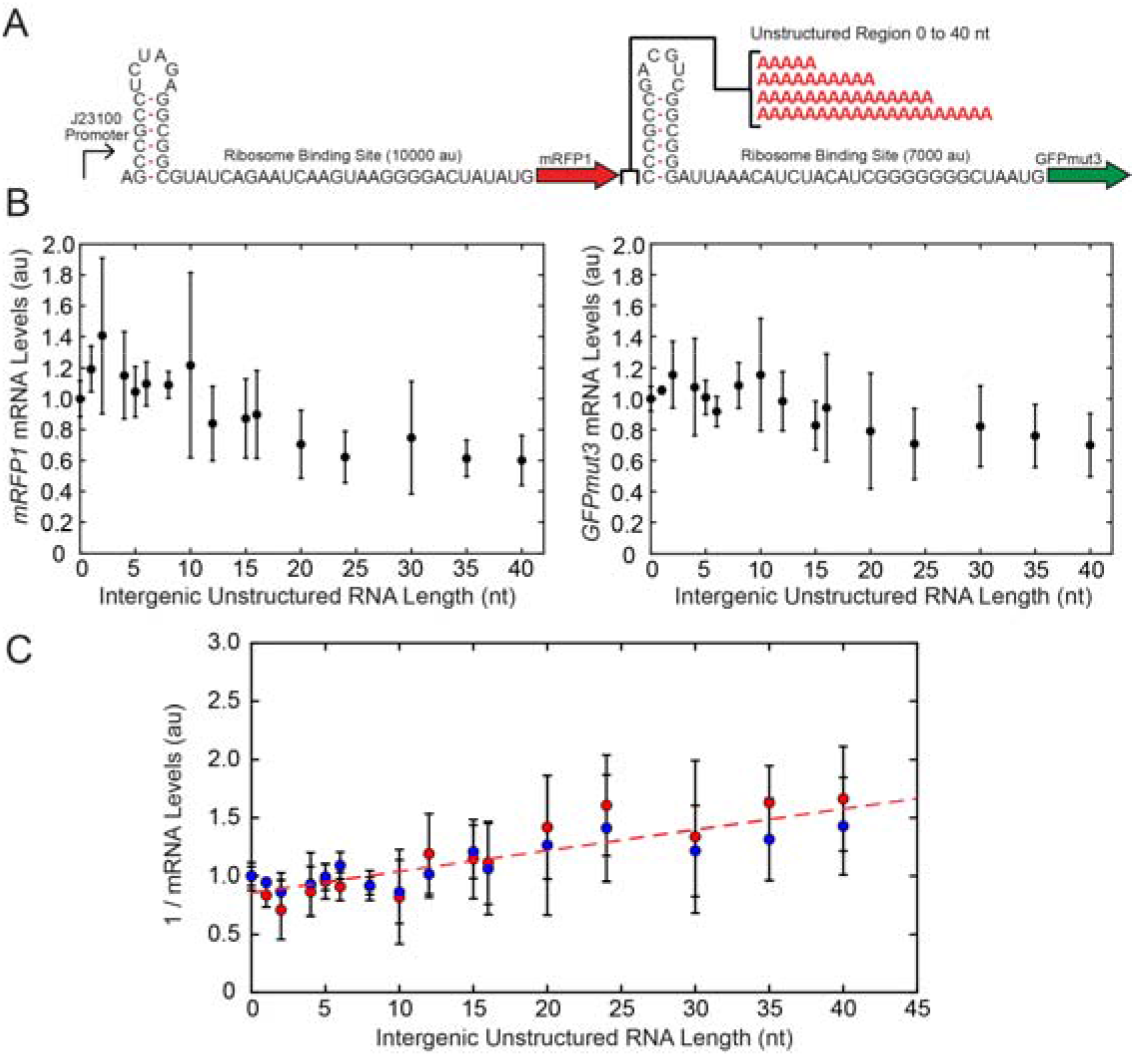
Sequence determinants of mRNA stability at intergenic sites. (A) Schematic illustrating bicistronic operons and the insertion of single-stranded RNA (RNAse E/G binding sites) with varying lengths into the intergenic region. (B) Characterization of the operons, showing (left) *mRFP1* mRNA levels compared to the single-stranded RNA lengths and (right) *GFPmut3* mRNA levels compared to the singlestranded RNA lengths. (C) The inverse mRNA levels for (red) *mRFP1* and (blue) *GFPmut3* are compared to the single-stranded RNA lengths alongside a (red dashed line) linear relationship with slope C = 0.018. Dots and error bars are the mean and standard deviation from 2 biological and 6 technical replicates.

As before, we replotted our measurements to illustrate how the inverse *mRFP1* and *GFPmut3* mRNA levels – proportional to the mRNA’s decay rate – are controlled by the ssRNA length within intergenic regions (**Figure 5C**). Similar to 5’ UTRs, we found that the ssRNA region must be at least 2 nt long before we observe any increase in mRNA decay. However, unlike 5’ UTRs, we observed only a linear quantitative relationship between ssRNA length and mRNA decay with a small slope (activity coefficient C = 0.018). Overall, these results show that RNAse E/G is not able to strongly bind and cleave intergenic regions even with large ssRNA regions, resulting in much smaller changes in mRNA decay rates. Moreover, we do not observe appreciable amounts of differential mRNA decay across upstream and downstream cistrons (indicative of polarity) or attenuation of mRNA decay at higher ssRNA lengths (indicative of transient loop formation).

### Transcriptional Terminator Efficiency Has No Effect on mRNA Stability

For our last dataset, we investigated how the efficiency of an operon’s transcriptional terminator affected its mRNA stability. To do this, we designed and constructed 8 operons expressing mRFP1, inserting different intrinsic transcriptional terminators into their 3’ UTRs (**Figure 6A**). Each terminator was selected from a toolbox of well-characterized synthetic terminators to have varying termination efficiencies from 8.3% to 99.7% (69). We also constructed a control operon that does not have a transcriptional terminator at the insertion position (0% efficiency). As an additional test, we also searched for any transcriptional terminator sequences downstream of the insertion site. Using the FindTerm terminator finder (70), the closest transcriptional terminator candidate was at least 230 base pairs downstream of the *mRFP1* stop codon. Otherwise, all operons have identical promoters, ribosome binding sites, and protein coding sequences.

**Figure 6:**
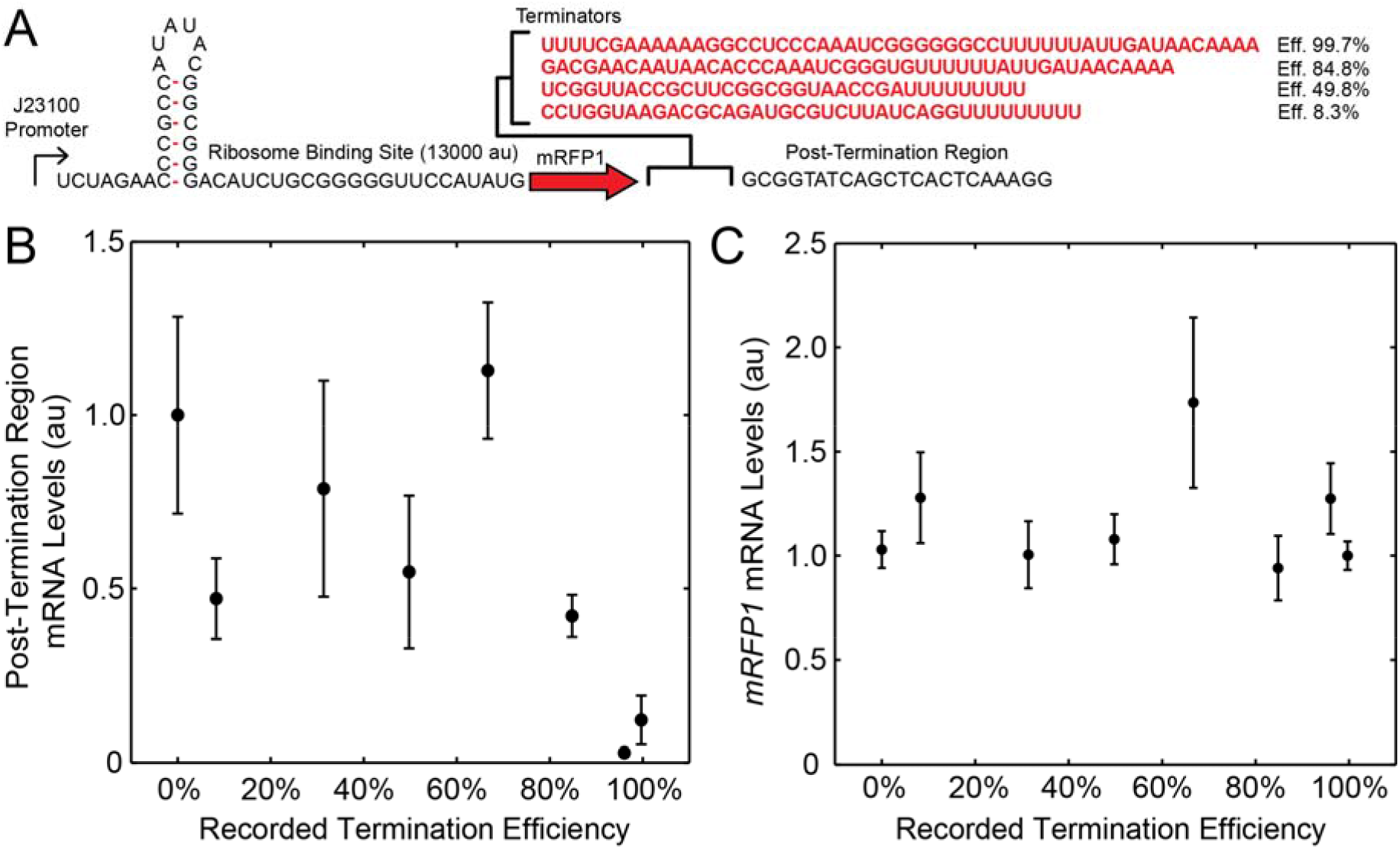
Transcriptional terminator efficiency does not impact mRNA stability. (A) Schematic illustrating *mRFP1* operons containing transcriptional terminators with varied efficiencies. (B) Characterization of the post-terminator mRNA levels using SYBR Green RT-qPCR assays, compared to prior measurements of the transcriptional terminators’ efficiencies. (C) Characterization of the *mRFP1* mRNA levels using TaqMan RT-qPCR assays as compared to the transcriptional terminators’ efficiencies. Dots and error bars are the mean and standard deviation from 2 biological and 6 technical replicates.

As before, we characterized the mRNA levels of these operons during exponential growth. For our first measurement, we applied SYBR Green RT-qPCR to measure the mRNA level of the region downstream of the inserted transcriptional terminators and to quantify the efficiencies of the transcriptional terminators (**Figure 6B**). We found that the levels of the post-terminator mRNA region were only greatly reduced once the previously recorded terminator efficiencies exceeded 80%. At the same time, we applied TaqMan RT-qPCR to measure the *mRFP1* mRNA levels. Surprisingly, we found no appreciable differences in *mRFP1* mRNA level across the full range of terminator efficiencies (**Figure 6C**). The *mRFP1* mRNA levels were similar whether a highly efficient terminator, a low efficiency terminator, or a non-terminating control sequence were inserted downstream of the *mRFP1* coding sequence. For the variants with little or no termination efficiency at the insertion site, it is likely that RNA polymerase continues to transcribe the mRNA for at least 230 nucleotides past the *mRFP1* stop codon, which would greatly lengthen the 3’ UTR. These results suggest that the length and structure of the 3’ UTR has little appreciable effect on the mRNA’s decay rate, which is consistent with the prior insertion of unstructured RNA into the 3’ UTR region (**Figure 4**).

## DISCUSSION

We designed and characterized 82 operons to quantify the sequence and structural determinants controlling mRNA stability in *E. coli*. We introduced rationally designed sequences into the 5’ UTR, intergenic, and 3’ UTR regions within mono-cistronic or bi-cistronic operons to measure how they affected the operons’ mRNA levels. Through this learn-by-design process, we systematically investigated how these determinants affected each mRNA decay pathway (**Figure 1**), including (i) the amount of unstructured, single-stranded RNA (ssRNA) in the 5’ UTR, which controls the rate of RNAse E/G cleavage via the end-dependent pathway (**Figure 2B**); (ii) the first three nucleotides of the transcript, which controls RppH-DapF dephosphorylation at the 5’ end (**Figure 2E**); (iii) the translation rate of coding sequences, which controls the amount of ribosome protection (**Figure 3**); (iv) the types and locations of RNAse binding sites, which controls decay rate and polarity (**Figure 4**); (iv) the amount of singlestranded RNA in the intergenic region, which controls the rate of RNAse E/G cleavage via the direct entry pathway (**Figure 5**); and (v) the efficiency of the transcriptional terminator, which controls the sequence, structure, and length of the 3’ UTR (**Figure 6**).

Overall, modifying the 5’ UTR of a mRNA transcript had the largest effect on its stability. Inserting a 20 nucleotide single-stranded RNA (ssRNA) region into the 5’ UTR led to a 9.4-fold decrease in mRNA level. Lowering the translation initiation rate of the first coding sequence by 35-fold – through redesigning its ribosome binding site – led to a 11.8-fold decrease in mRNA level. Modifying just the first three nucleotides of the transcript could lead to 18-fold change in mRNA level! In contrast, changing the intergenic region’s sequence and structure had much less potent effects. Inserting a 20 nucleotide ssRNA region into the intergenic region reduced mRNA levels by up to 2-fold, similarly affecting both upstream and downstream genes in a bi-cistronic operon (no polarity). Inserting a long RNA hairpin into the intergenic region mainly affected the downstream coding sequence’s mRNA level (by 1.8-fold). Finally, we were genuinely surprised that our substantial changes to the 3’ UTR sequence, structure, and length had no appreciable effect on the transcript’s mRNA level. Based on our measurements, the transcriptional terminator does not alter a mRNA’s decay rate and only facilitates decoupling of cistrons in adjacent operons. Overall, our results show that end-dependent decay – mediated by RNAse E/G and RppH-DapF – is the predominant pathway controlling mRNA stability. RNAse E/G likely bind internal unstructured sites with much lower affinity, requiring long stretches of unstructured RNA (e.g. entire coding regions) to achieve high cleavage rates.

Leveraging our measurements, we developed several biophysical models to explain how these sequence determinants controlled mRNA decay rates. Each model contains a small number of parameters, but can explain the observed quantitative trends. Previously, RNAse E/G were suggested to bind to specific binding motifs (e.g. RAUUW or RNWUU). Here, we propose that RNAse E/G simply bind unstructured RNA regions with a 2 nucleotide landing pad. Longer unstructured regions therefore provide a proportionally larger number of potential binding sites for RNAse E/G with an expected increase in cleavage rate (**Equation 1**). When this model is applied to the 5’ UTR, it fits our mRNA decay rate measurements extremely well for when the ssRNA length is varied from 1 to 20 nucleotides (**Figure 2C**). However, when the 5’ UTR ssRNA region is very long, we see an attenuation of this effect, suggesting that the region is no longer fully unstructured. We therefore turned to polymer theory, which provides well-established equations for calculating the likelihood that a polymer forms transient, non-specific structures (ie. wormlike chains), which determines the fraction of polymer that remains accessible. By incorporating polymer theory (**Equation 2**), a single model was able to explain how ssRNA length affects mRNA decay rates across the larger range of lengths from 1 to 40 nucleotides, using a persistence length that is consistent with prior measurements.

We applied the same biophysical modeling (**Equation 1**) to the intergenic region with a similarly consistent explanation. Longer ssRNA regions led to proportionally higher mRNA decay rates (**Figure 5**). However, the proportionality constant in this model was much lower, consistent with our observations that the direct entry pathway is much slower than the end-dependent pathway, when acting on the same amount of ssRNA. Notably, unlike 5’ UTRs, intergenic regions with large amounts of ssRNA had the highest amounts of mRNA decay, which is expected because the formation of transient wormlike structures requires at least one freely movable (untethered) end. While the 5’ end of the transcript can freely move, both ends of the intergenic region are physically constrained by the surrounding ribosomes engaged in translation elongation.

We also developed a biophysical model of ribosome protection that shows how systematically varying a coding sequence’s translation rate controls its mRNA level (**Figure 3**), applying equations to calculate a mRNA transcript’s ribosome density and ribosome-to-ribosome headway distance (**Equations 3–6**). The model provides a mechanistic basis for the observed sigmoidal-like relationship in our measurements with expected plateaus in mRNA level at either very low or very high translation rates, and with sharp changes in between. Here, we treated the CDS region as being equally accessible to RNAse E/G activity regardless of their RNA structure. We also assumed that all codons have the same translation elongation rates. In return, the resulting equations can be analytically solved and remain remarkably simple to evaluate. However, the observed differences in proportionality constants for *mRFP1* and *sfGFP* could arise from these assumptions.

Collectively, our biophysical models suggest straight-forward rules for designing operons with controlled mRNA stabilities. For maximum stability, 5’ UTRs should begin with 5’-AGN ends and support moderate-to-high translation initiation rates (10000 or higher on the RBS Calculator v2.1 scale), while limiting the amount of single-stranded RNA outside the ribosome binding site. For multi-cistronic operons, intergenic regions should also contain ribosome binding sites that support moderate-to-high translation initiation rates, though the amount of single-stranded RNA will not appreciably affect their stability. Our design rules here do not consider the presence of self-cleaving ribozymes, which introduce a 5-hydroxyl group that slows down end-dependent mRNA decay (27). To achieve lower mRNA stabilities, transcripts can be designed to have 5’-AAN ends as well as single-stranded RNA regions in their 5’ UTR regions, upstream of the ribosome binding site. Perhaps most surprisingly, the design of the 3’ UTR does not affect the operon’s expression levels and primarily controls read-through transcription into downstream operons. Altogether, our measurements and models provide the quantitative means to controlling mRNA stability in operons.

## SUPPLEMENTARY DATA

Supplementary Data are available at NAR online.

## CONTRIBUTIONS

D.P.C. and H.M.S designed the study, analyzed results, and wrote the manuscript. D.P.C. conducted the experiments.

